# ROSeq: Modeling expression ranks for noise-tolerant differential expression analysis of scRNA-Seq data

**DOI:** 10.1101/374025

**Authors:** Krishan Gupta, Manan Lalit, Aditya Biswas, Ujjwal Maulik, Sanghamitra Bandyopadhyay, Gaurav Ahuja, Abhik Ghosh, Debarka Sengupta

**Affiliations:** Department of Computer Science and Engineering, Indraprastha Institute of Information Technology, New Delhi, 110020, India; Max Planck Institute of Molecular Cell Biology and Genetics, Dresden 01307, Germany; Microsoft India Pvt. Ltd., Hyderabad, Telangana 500032, India; Department of Computer Science, Jadavpur University, Kolkata, West Bengal 700032, India; Machine Intelligence Unit, Indian Statistical Institute, Kolkata 700108, India; Department of Computational Biology, Indraprastha Institute of Information Technology, New Delhi, 110020, India; Interdisciplinary Statistical Research Unit, Indian Statistical Institute, Kolkata 700108, India

## Abstract

Systematic delineation of complex biological systems is an ever-challenging and resource-intensive process. Single cell transcriptomics allows us to study cell-to-cell variability in complex tissues at an unprecedented resolution. Accurate modeling of gene expression plays a critical role in the statistical determination of tissue-specific gene expression patterns. In the past few years, considerable efforts have been made to identify appropriate parametric models for single cell expression data. The zero-inflated version of Poisson/Negative Binomial and Log-Normal distributions have emerged as the most popular alternatives due to their ability to accommodate high dropout rates, as commonly observed in single cell data. While the majority of the parametric approaches directly model expression estimates, we explore the potential of modeling expression-ranks, as robust surrogates for transcript abundance. Here we examined the performance of the Discrete Generalized Beta Distribution (DGBD) on real data and devised a Wald-type test for comparing gene expression across two phenotypically divergent groups of single cells. We performed a comprehensive assessment of the proposed method, to understand its advantages as compared to some of the existing best practice approaches. Besides striking a reasonable balance between Type 1 and Type 2 errors, we concluded that ROSeq, the proposed differential expression test is exceptionally robust to expression noise and scales rapidly with increasing sample size. For wider dissemination and adoption of the method, we created an R package called ROSeq, and made it available on the Bioconductor platform.

## 2 Introduction

In the past few years, single cell RNA-Sequencing (scRNA-Seq) has dramatically accelerated the characterization of molecular heterogeneity in healthy and diseased tissue samples [17]. The declining cost of library preparation and sequencing have fostered the adoption of single cell transcriptomics as a routine assay in studies arising from diverse domains, including stem cell research, oncology, and developmental biology [23, 6]. The field of single cell transcriptomics suffers severely from various data quality issues, mainly due to the lack of starting RNA material. High levels of noise and technical bias pose significant challenges to single cell gene expression modeling, which hinders arrival at statistically apt conclusions about cell-type-specific gene expression patterns [14].

A number of parametric and nonparametric methods have already been proposed for modeling single cell expression data and finding differentially expressed genes (DEGs). SCDE [5], MAST [2] and BPSC [21] are notable among these. SCDE and MAST model gene expression using well-known probability density functions and mixture models involving some of those. BPSC, on the other hand, handles single cell expression bimodality by employing a Beta-Poisson mixture. Different from these, we conjectured that considering expression ranks instead of absolute expression estimates would make a model less susceptible to the noise and the technical bias, as commonly observed in single cell data. To realize the same, we employed Discrete Generalized Beta Distribution (DGBD) [11] to model the distribution of expression ranks instead of the raw count. The consideration of rank-ordering distribution was inspired by the seminal work by Martinez-Mekler and colleagues, where they demonstrated the universal applicability of the same in linking frequency estimates and their ranks [11]. We developed ROSeq, a Wald-type test to determine differential expression from scRNA-Seq data.

On real data, we demonstrated that DGBD efficiently approximates cell lineage-specific marginal distribution of gene expression levels. Further, to explore if our approach of modeling expression ranks performs competitively in identifying differentially expressed genes, we performed a series of experiments involving simulations and validation against ground truth information. We assessed the robustness of differential expression calls by injecting artificial noise into real data. In line with our expectation, ROSeq was found to be the least perturbed by noise-driven distortions of the expression signal.

## 3 MATERIALS AND METHODS

### 3.1 Data description

We used three publicly available single cell RNA sequencing (scRNA-Seq) datasets for the various analyses. For better readability, we name the datasets after the first authors’ surnames. Among these, the Trapnell data contains scRNA-Seq profiles of 77/99 primary myoblasts sampled before/24 hours after differentiation [18]. Tung data consists of single cell transcriptomes of induced pluripotent stem cells (iPSCs) generated from three different individuals, marked as NA19098, NA19101, and NA19239, respectively [19]. A total of 288 cells were profiled for each of the three individuals. For both the Trapnell and the Tung datasets, three bulk RNA-Seq replicates were available from the respective studies for each condition/individual. Notably, both Trapnell, as well as Tung datasets, were generated using the SMARTer chemistry. To diversify our experiments, we used the Zheng dataset containing 3258 single cell transcriptomes of Jurkat cells, processed using the GemCode technology [22].

### 3.2 Data Pre-processing

For each dataset, we first filtered out cells having less than 2000 detected (non zero read count) genes. We retained having read count > 3 in at least 3 cells [4]. The processed count matrix was subjected to different normalization techniques depending on the target differential expression method. For Wilcoxon’s rank-sum test, BPSC, and MAST, count per million (CPM) normalization was used, following the recommendation by Soneson and Robinson [16]. SCDE and DESeq2 [9] received the processed raw count data as input. For ROSeq, we first subjected the processed raw count matrices to the trimmed mean of M-values (TMM) normalization [13], followed by Voom transformation [8].

### 3.3 Mapping expression estimates to ranks

For gene expression modeling, ROSeq accepts normalized read count data as input. For each gene, ROSeq first defines its range by identifying the minimum and the maximum values by pulling the normalized expression estimates across both the cell-groups under study. Next, the range is split into *k* × σ sized bins, where *k* is a scalar with a default value of 0.05, and σ is the standard deviation of the pulled expression estimates across the cell-groups. Each of these bins is assigned a rank, based on the sequential order of its expression range. At the level of a cell-group, this leads to mapping of bin-wise cell frequencies to ranks, such that the bin with the highest cellular frequency is assigned the least rank (i.e., 1). The Discrete Generalized Beta Distribution (DGBD) is used as a probability mass function to express a normalized bin-wise cell-frequency *y*_*r*_ as a function of its corresponding rank *r* using two real parameters *a* and *b*. Let *N* be the total number of bins for a given gene. Then the DGBD formulation is expressed as

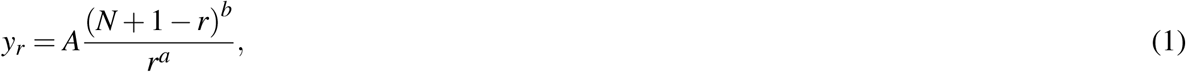

where *A* is the normalizing constant ensuring that the sum of the normalized frequencies equals one.

### 3.4 Estimation of the DGBD parameters

For a given gene and a specific cell-group, the best-fitting parameter values 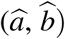 are determined by maximizing, with respect to (*a, b*), the Log-Likelihood corresponding to the model given by Equation 1. From the DGBD expression, the required Log-Likelihood function, **logL**, can be computed as (in Equation 2)

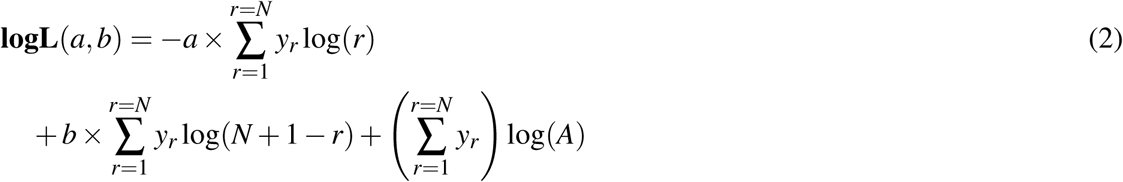

The resulting estimates 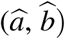 correspond to the DGBD under which the observed data is most likely to be generated. Such maximum likelihood estimates (MLE) are the most efficient (least standard error) and enjoy several optimum properties on large sample sizes [1].

To test differential expression of a gene between two cell-groups, based on the above MLEs 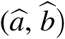, we additionally need estimates of their standard errors (equivalently their variance). From the theory of maximum likelihood [12], the asymptotic variance of 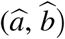 is given by the inverse of the associated Fisher information matrix *I*(*a, b*)^−1^, which can be consistently estimated by 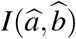. For the log-likelihood function of the DGBD model given in Equation 2, the form of the Fisher information matrix *I* may be simplified in a more succinct form as follows.

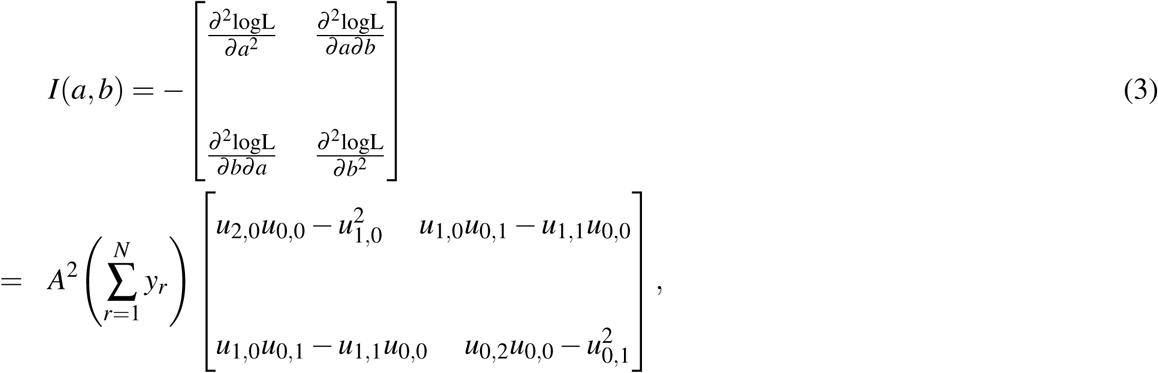

where, for each *i, j* = 0, 1, 2, we define

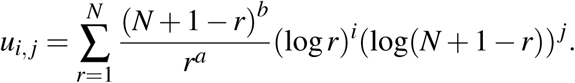

Note that, *u*_0,0_ = 1*/A*. See the Supplementary Note for the derivation of *I*(*a, b*).

### 3.5 Testing for differential expression: Two-sample Wald Test

Further, in order to statistically test if a gene is differentially expressed between two sub-populations, ROSeq uses the (asymptotically) optimum two-sample Wald test based on the MLE of the parameters and their asymptotic variances, given by the inverse of the Fisher information matrix.

Let us assume that the DGBD parameters corresponding to the contrasting cell-groups 1 & 2 are denoted by (*a*_1_, *b*_1_) and (*a*_2_, *b*_2_), respectively, and their MLEs based on the available normalized expression data are given by 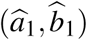 and 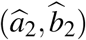 with the respective number of bins being *m* and *n*. We can estimate the asymptotic variance matrices for these MLEs, using Equation 3, as 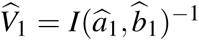 and 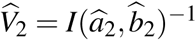, respectively. Under our the DGBD model, the desired testing for differential gene expressions is equivalent to the test for the null hypothesis *H*_0_ : *a*_1_ = *a*_2_, *b*_1_ = *b*_2_ against the omnibus alternative. The Wald test statistic *T* for testing *H*_0_ can be written as follows:

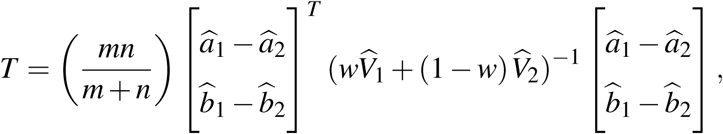

where 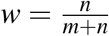. If the null hypothesis *H*_0_ is correct, i.e., the genes in the two sub-populations are not differentially expressed, the above test statistics *T* follows a central chi-square distribution 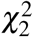 with two degrees of freedom. Therefore, we conclude that the genes are differentially expressed (i.e., reject *H*_0_) at 95% level of significance, if the observed value of the test statistics *T* exceeds the 95% quantile of the 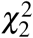 distribution (which is approximately 6). The corresponding *P*-value is given by the probability that a 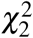 random variable exceeds the observed value of *T*.

### 3.6 Introducing noise to expression data

To assess the robustness of various DEG callers, we introduced noise to gene expression values of Jurkat cells from the Zheng dataset and evaluated the Type I error control corresponding to each of the methods. To add noise the following steps were implemented — 1. Null datasets were constructed by randomly selecting Jurkat cell transcriptomes into two equal sized groups. The number of cells in the groups varied from 25 to 250, with a step-wise increase of 25. 2. Next, for each gene, one of the cell-groups was chosen, and 10% of the read count values (across cells in the concerned group) were added with simulated Gaussian noise. For a read count value *r* under alteration, the updated read count value *r*^′^ was set to *r* + *𝒩* (*m* + *c.s, k.s*), where *m* is the mean and *s* is the sample standard deviation of the read count values for the concerned gene across cells of the chosen group. Notably *c* and *k* are constants. We experimented with *c* = 2, 4 and *k* = 0.2, 0.4.

## 4 Results and Discussion

### 4.1 Overview of ROSeq

Advanced droplet-based single cell RNA sequencing technologies can profile several thousands of cells in a single experiment [10, 22]. Despite considerable advancements, expression readouts obtained from these high-throughput platforms suffer from various technical and trivial biological distortions [20]. These include single cell library size differences, cell cycle effects, amplification bias, low RNA capture rate, and high levels of dropout events [5]. Different from bulk RNA sequencing, gene expression modeling in single cells requires special statistical considerations [3]. Since the introduction of the single cell technologies, numerous parametric models have been proposed, primarily accounting for dropout events. The majority of these are mixture models of distinct probability density functions. We conjecture that a limitation of such approaches could be that they disregard the other noise sources as enumerated above. Ranks are commonly known to be more robust as compared to the corresponding expression estimates. In fact, with the increase in sample size, single cell studies are now seen embracing the traditional Wilcoxon’s rank-sum test to identify differentially expressed genes. While non-parametric methods are assumption-free [14], they often lack statistical power. In this work, we explored the utility of discretizing an expression vector into bins and ordering them (meaning ordering ranks corresponding to bins) based on bin-wise cellular frequencies, thereby making it modellable by Discrete Generalized Beta Distribution (DGBD) (aka, rank-ordered distribution [11]). Notably, fitting DGBD involves MLE of two shape parameters, denoted by *a* and *b*. Figure 1 depicts an example of DGBD based modeling of COP9 Signalosome Subunit 6 (*COPS6*) expression across 288 single cells from the biological replicate NA19098 of the Tung data [19]. For a comprehensive assessment of the quality of fit, we estimated *R*^2^ for all the 11513 genes that qualified the filtering criteria. Quite remarkably, DGBD fits yielded *R*^2^ > 0.9 for a vast majority of the genes, thereby underscoring its appropriateness in modeling expression ranks.

**Figure 1.**
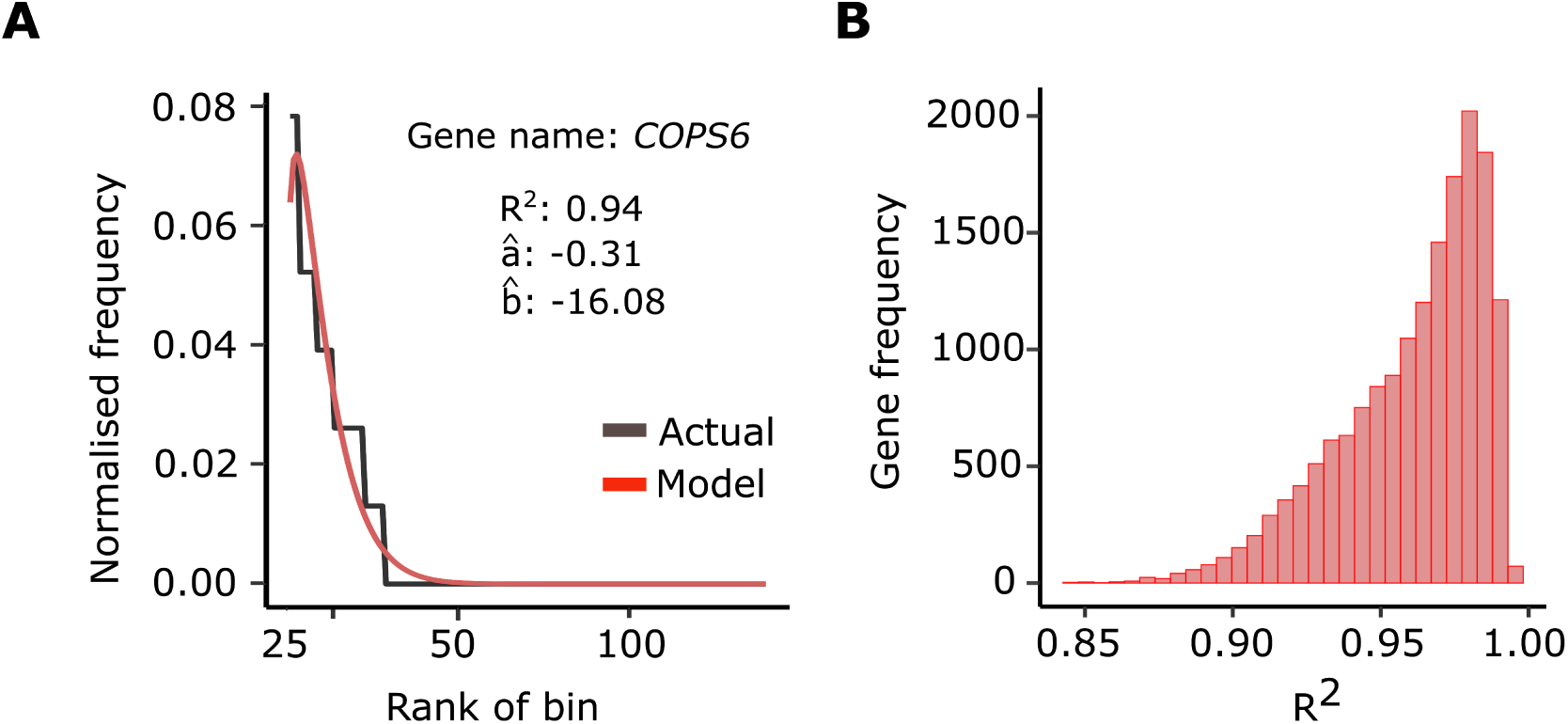
Modeling xssingle cell expression data. (A) Discrete Generalized Beta Distribution (DGBD) based modeling of *COPS6* expression. Discretized expression bins are ranked based on normalised bin-wise cellular frequencies. (B) Distribution of *R*^2^ values obtained from DGBD based modeling of 11513 expressed genes (Tung data).

### 4.2 Rate of Type I errors

Leveraging DGBD based modeling of expression, we devised ROSeq, a Wald type test to determine differential expression in single cell data. To evaluate Type I error control associated with ROSeq, we constructed several null datasets, by segregating cells of the same type into two groups, for varied group sizes [16]. For each of the methods, we tracked the fraction of the tested genes that were assigned a nominal adjusted *P*-value of less than 0.5. We iterated this simulation experiment for varied cell-group sizes — 25, 50, 75, 100, 125, and 150. For each cell-group size, 100 null datasets were constructed and subjected to the various DEG callers. For this experiment we used single cell expression profiles of iPSCs (replicate id: NA19098) from the Tung dataset [19]. In addition to ROSeq, five other methods, namely BPSC, SCDE, Wilcoxon’s rank-sum test, MAST, and DESeq2, were considered for performance comparison. Among all the six methods, SCDE offered the overall best performance on small-sized cell groups, closely matched by ROSeq, with an increase in sample size. Among the remaining methods, MAST performed reasonably well, followed by DESeq2, whereas Wilcoxon and BPSC faired poorly with comparable Type I error rates (Figure 2). Notably, on small sample sizes, ROSeq’s performance remains the worst, which betters steadily with the increase in the number of cells. Conversely, SCDE exhibits extreme conservativeness with minimal false discovery rates, irrespective of the sample size.

**Figure 2.**
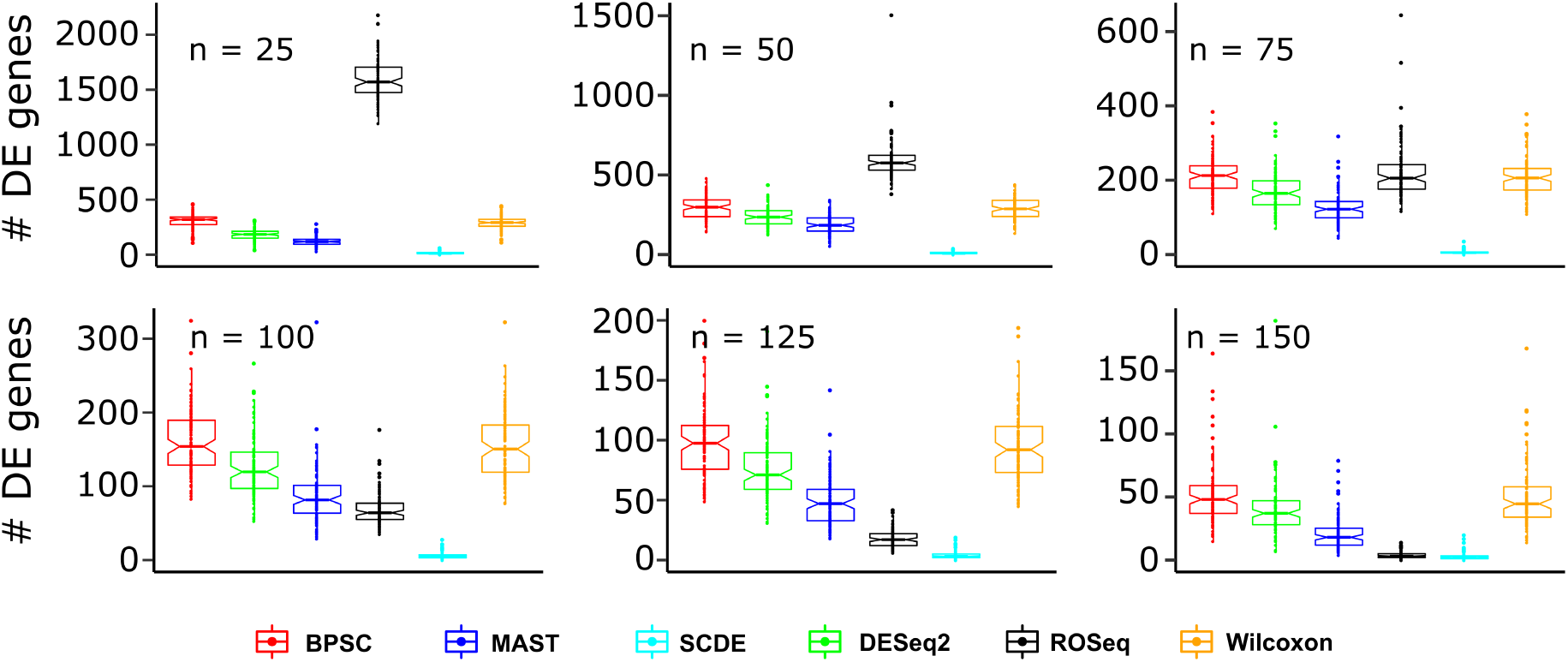
Boxplots showing Type I error rates obtained by applying different DEG callers on null datasets, for varied cell-group sizes. For each considered group-wise cell count, 100 experiments were performed to track the variability in Type I error rates. The associated distributions are depicted by the boxplots.

### 4.3 Comparative benchmarking based on matched bulk RNA sequencing data

Tissue-level measurement of gene expression is considered more robust compared to single cell-based estimates. As such, it’s a common practice to benchmark DEG calls on single cells w.r.t. DEGs obtained from matched bulk expression profiles. We accessed scRNA-Seq data from two previous studies that also performed bulk RNA-Seq on the same samples. Description of the datasets can be found in the Materials and Methods section. Four contrasting cell-group pairs were constructed as follows — myoblasts before and 24 hours after differentiation (source: Trapnell data), and all three combinations of biological replicates of human iPSCs (source: Tung data). For both the datasets, three technical replicates were available, which we used for confident DEG calls, using DESeq2 (FDR cutoff of 0.05). Single cell DEG calls were made using all six methods, including ROSeq as described previously. A single cell DEG call was considered true positive if the gene was also present in the matched bulk-transcriptome DE list. If not, it was counted as a false positive. ROC-AUC values were computed for an objective comparison of the methods. Out of four datasets, on three, ROSeq and SCDE topped among all the methods with negligible difference in ROC-AUC values Figure 3. For one of the three cases associated with the Tung data, SCDE marginally outperformed ROSeq. Although DESeq2 is not specialized for single cells, we used it as a control to ensure single cell focused methods yield overall better performance. Notably, on three out of the four datasets, Wilcoxon’s rank-sum test struggled to match ROSeq’s performance.

**Figure 3.**
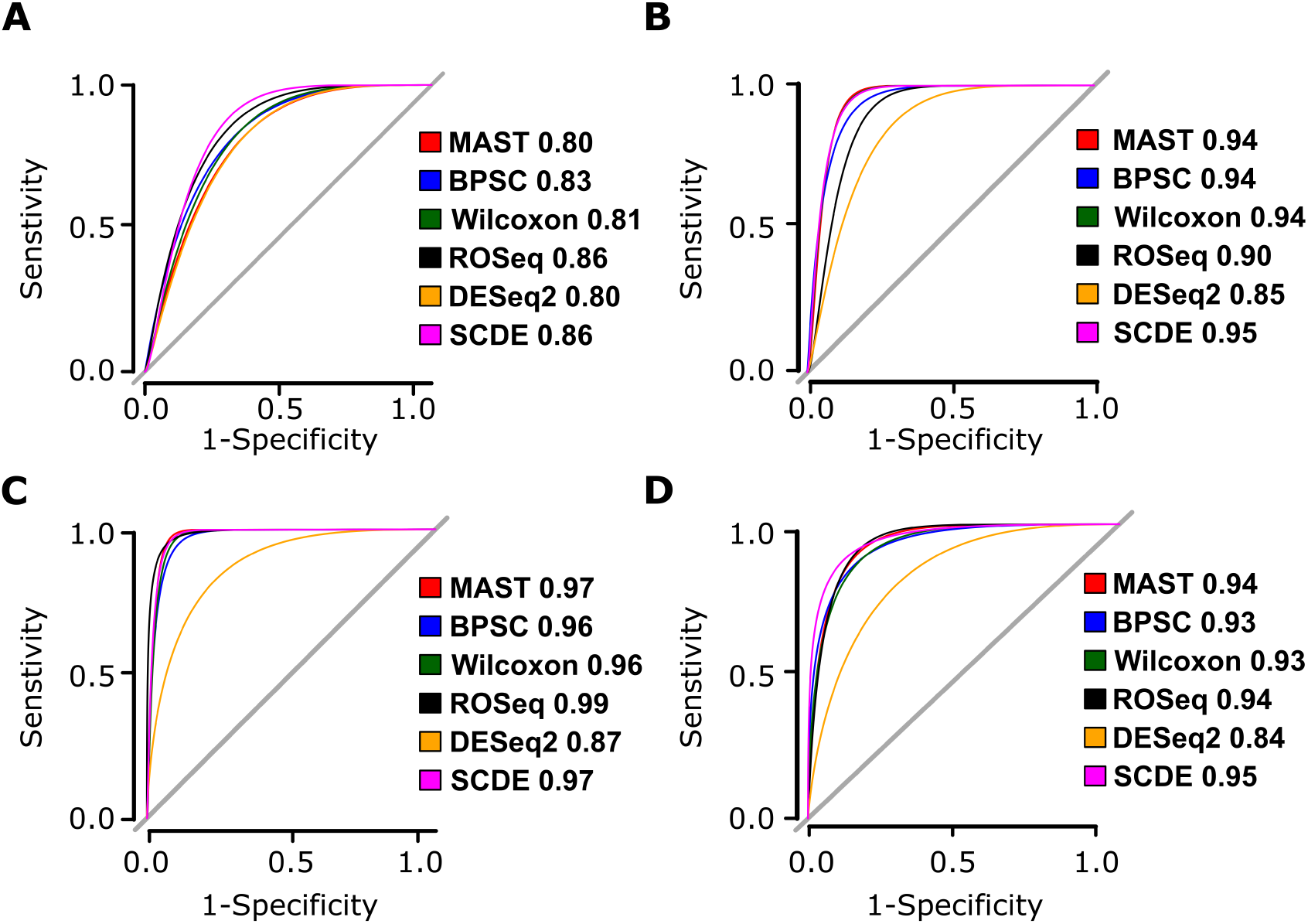
Bulk RNA-Seq based benchmarking of DEG calls. (A) Receiver Operating Characteristics curve (ROC) and the associated Area Under the Curve (AUC) values obtained by bulk-based benchmarking of single cell DEG calls between myoblsats at 0 and 24 hours (Trapnell data). (B) Similar analysis between iPSCs collected from replicates NA19098 and NA19101 (Tung data), (C) replicates NA19098, and NA19239 (Tung data), (D) replicates NA19101, and NA19239.

Besides ROC-AUC, we also tracked other popular measurements of classification accuracy, including Mathews Correlation Coefficient (MCC) [15], and Cohen’s Kappa (*k*) [7], which control for random performance. MCC and *k* are more reliable than ROC-AUC, when the involved categories (in this case, true positive and true negative DEGs) are not balanced (Supplementary Table). Notably, for all three cases associated with Tung data, ROSeq maximized the MCC and *k* scores, whereas in the case of Trapnell data, SCDE outperformed ROSeq. We predict, subpar MCC and *k* scores in case of Trapnell data could be linked with the lack of cells in the concerned cell-groups, as realized while evaluating the Type I error control.

Upon viewing the expression distribution and rank-ordered distribution corresponding to some example DE/non-DE genes called by ROSeq, as per expectation, we noted that the latter significantly stabilizes the shape diversity, as observed in marginal distribution of expressed genes (Supplementary Figures 1, 2).

### 4.4 Noise tolerance of the DEG callers

As stated earlier, single cell transcriptomes are distorted by technical biases such as RNA degradation during cell isolation and processing, variable reagent amounts, presence of cellular debris, and PCR amplification bias. Further, due to the small number of detected molecules, single cell expression estimates are inherently noisy, even in the absence of technical variability [14]. In a controlled experiment, we compared the DEG callers for their noise tolerance ability, measured in terms of false-positive DEG calls on null data, created by sampling Jurkat transcriptomes from the Zheng data [22]. For each gene, we introduced Gaussian noise to different sub-fractions of cells (10% and 20%) from one of the two cell-groups chosen randomly. ROSeq showed remarkable resilience to noise, outwitting the remaining methods by large margins (Figure 4 and Supplementary Figures 3). MAST and Wilcoxon’s rank-sum test performed reasonably well, whereas the remaining methods, i.e., DESeq2, BPSC, and SCDE, struggled to control false-positive DEG calls. We did similar experiments by injecting read count noise following uniform distribution. SCDE was found to be extremely robust in this case, neared by ROSeq with increase in cell numbers (Supplementary Figures 4, 5).

**Figure 4.**
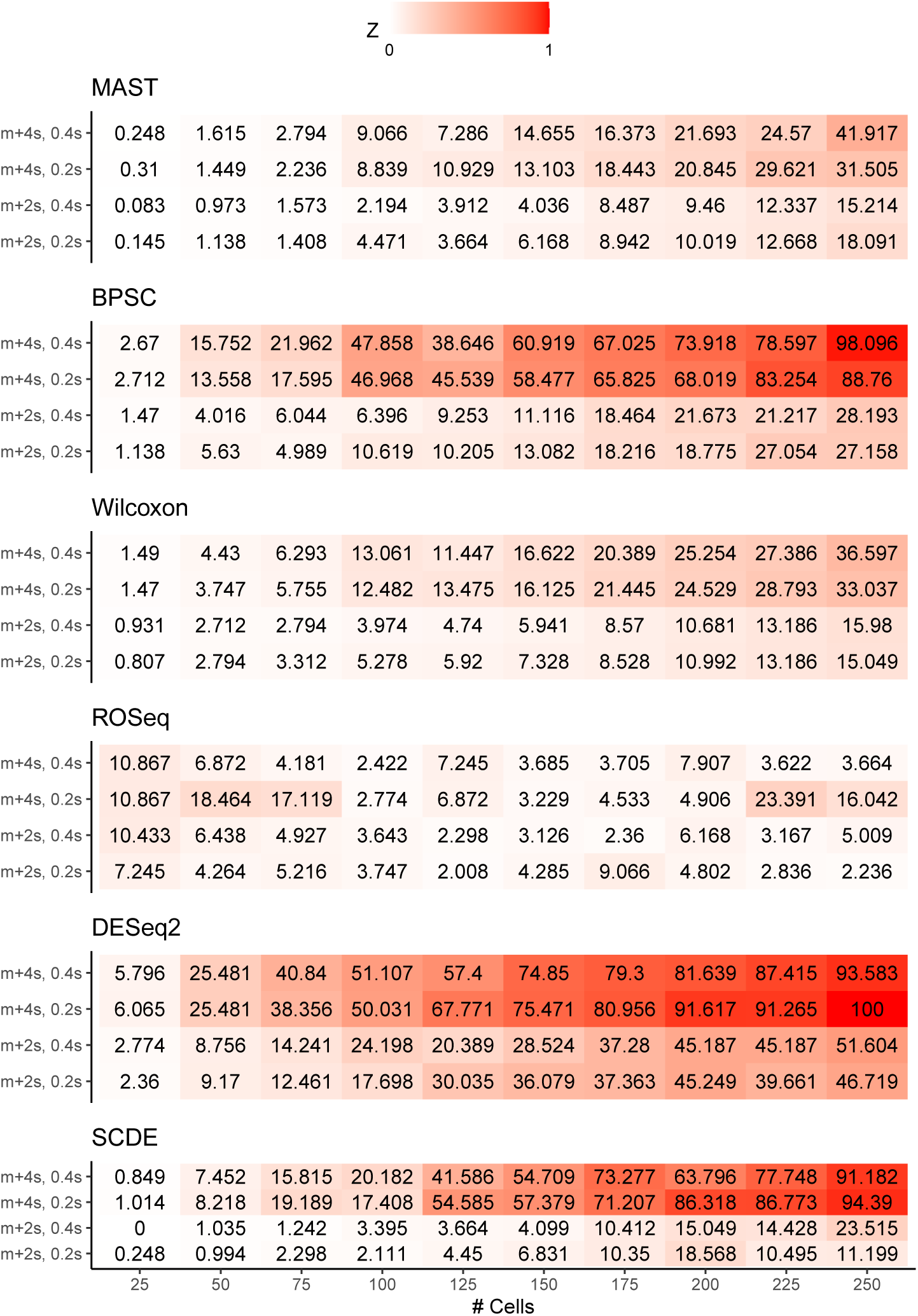
Analysis of noise tolerance. Each heatmap corresponds to a DEG caller, and signifies the Type I error rates under various statistical considerations. The X axis labels of the heatmaps indicate cell-count in each of the contrasting group of the null datasets, whereas the labels of Y axis indicate mean-standard deviation pairs corresponding to the Gaussian noise. The terms *m* and *s* indicate sample mean and standard deviation respectively derived from group specific read counts for the the genes under randomised alteration

### 4.5 Runtime efficiency

With the advent of droplet-based commercial platforms, profiling of tens of thousands of cells in a single experiment has become a common affair. Unsupervised clustering of large scale scRNA-Seq data produces numerous clusters with a significant number of cells in each of it. As such, besides accuracy, the scalability has become a desirable feature for the DEG callers. We, therefore, benchmarked time consumption by the methods for variable sizes of input scRNA-Seq datasets. For the construction of the datasets, we performed the same steps as we did for estimating the Type I error rates. The cell number in each group was varied between 10 and 150. As depicted by Figure 5, among all the six methods, Wilcoxon’s rank-sum test consumed the least time, followed by ROSeq/MAST. Although DESeq2, scaled fast for cell group sizes below 75. Post that DESeq2 turned slow compared to ROSeq/MAST. SCDE, remained the slowest, followed by BPSC. Notably, at log-scale, the order of growth in time consumption by ROSeq, MAST, and BPSC remained considerably flat, as compared to the remaining methods. Going by the growth-rate trends, we predict, ROSeq to be equally fast/faster than Wilcoxon’s rank-sum test. Notably, all experiments reported in this article were performed on a workstation configured with AMD Ryzen 7 3700X 8-Core processor with a clock speed of 4249.648 MHz, 64GB DDR4 RAM and Ubuntu 18.04.4 LTS operating system with 5.3.0-40-generic kernel. For fair bench-marking the time taken by each method was tracked by running it on a single core.

**Figure 5.**
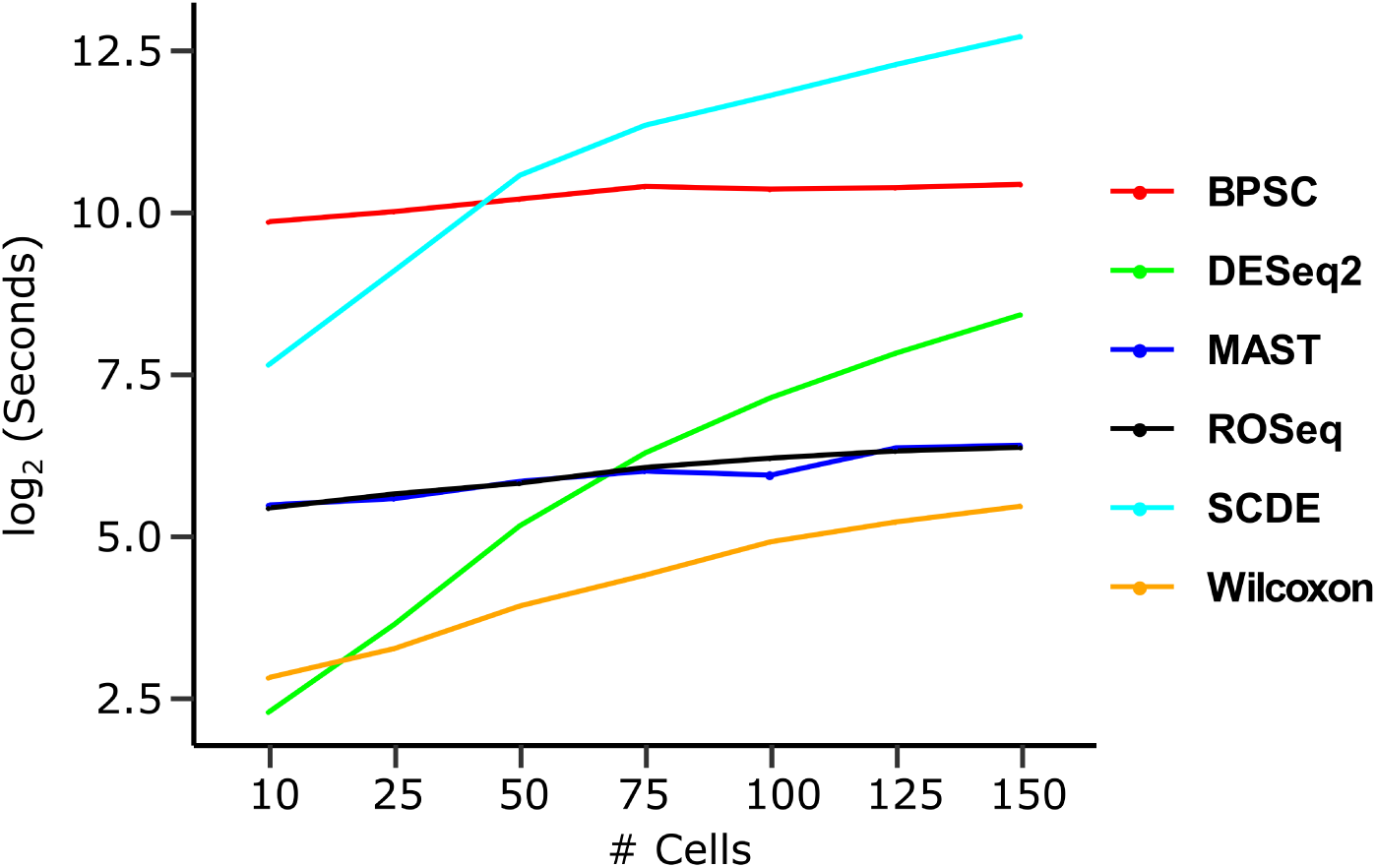
Time consumption by different methods as per group cell-count increases.

## 5 Conclusion

Martinez-Mekler and colleagues demonstrated that two-parameter DGBD (rank-ordered distribution) gives excellent fits to diverse phenomena, arising from the arts, social and natural sciences [11]. We evaluated the applicability of DGBD to gene expression data. We further developed ROSeq, a DGBD-based Wald type test for differential expression analysis of scRNA-Seq data. Most of the statistical models for single cell expression data use mixed models to accommodate high dropout rates. ROSeq discretizes the data, thereby stabilizing local distortions in the shape of the distribution, due to noise and technical bias. We found DGBD to fit well to the entire spectrum of expressed genes of varying expression levels.

We systematically compared the performance of ROSeq with some of the existing best-practice methods such as SCDE, MAST, BPSC, which are largely tailored for single cell expression data. Among various critical observations, our systematic tracking of Type I errors revealed that a relatively higher number of cells (at least 100 in each contrasting group) are required for ROSeq to attain optimal performance, comparable to SCDE. Current studies report hundreds to thousands of cells per unsupervised cell cluster with the advent of droplet-based single cell profiling platforms. As such, we do not foresee any hindrance to ROSeq’s applicability due to cell paucity. However, ROSeq might produce sub-optimal DEG calls if a cluster contains a small number of cells. This shortcoming can be attributed to ROSeq’s use of asymptotic distribution.

Statistical test of differential expression involves comparing the marginal distribution of a gene’s expression across two cell-groups. Conversely, ROSeq analyzes the distribution of rank-ordered discretized expression bins across two cell-populations. This could give rise to false-negative DEG calls in rare scenarios. For example, if cell-population wise expression distributions of a gene are identically shaped yet distantly located, ROSeq might fail to recognize its DEG status.

Benchmarking bulk tissue-based DEG calls underscored competitive performance by the methods, tailored for single cell expression data. However, SCDE and ROSeq maximized the DEG call accuracy. ROSeq is particularly powerful when the single cell based expression estimates are inherently noisy. In such cases, ROSeq by far outwits the other tested methods. Further, ROSeq scales comfortably with an increase in number profiled cells, thereby making it a promising alternative to the existing DEG callers.

## 6 AVAILABILITY

The ROSeq R package is available at the Bioconductor portal: http://www.bioconductor.org. A more frequently updated version of the software can be accessed at: Github.

## Supplementary Data

## 1 Supplementary Note: Derivation of the Fisher Information Matrix

Note that, for our DGBD model with likelihood function given by Eq (2) of the main paper, we have

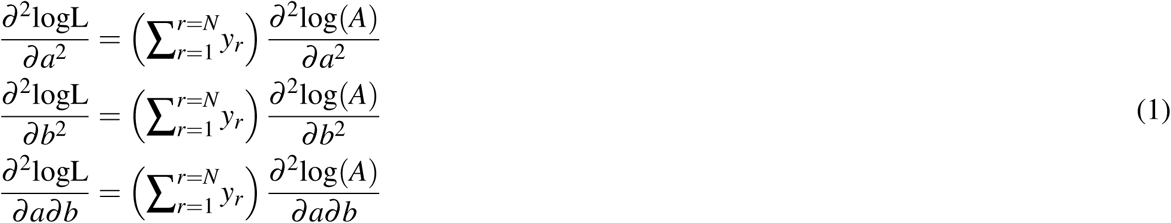

So in order to evaluate the above mentioned double derivatives, the first order derivative 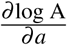 and 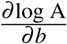 are determined as follows:

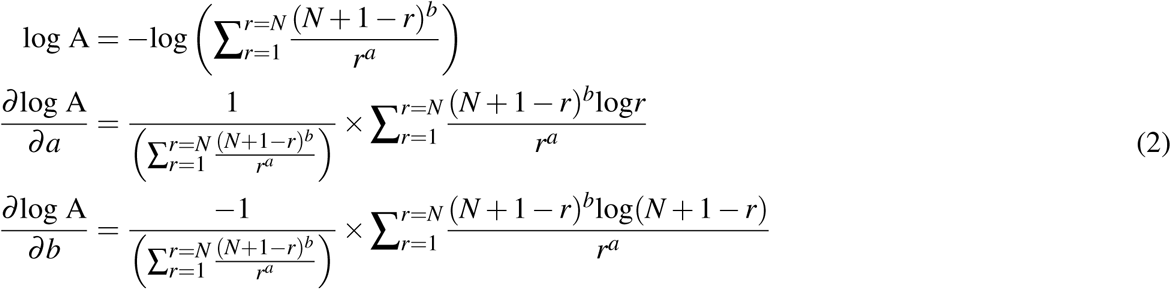

Re-writing Equation 2 in a more succinct form in the Equation 3 below, we get

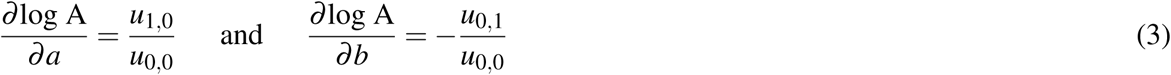

where *u*_*i, j*_s are as defined in the main paper. Evaluating the partial derivatives of *u*_1,0_, *u*_0,0_ and *u*_0,1_ with respect to *a* and *b*, in the Equation 4:

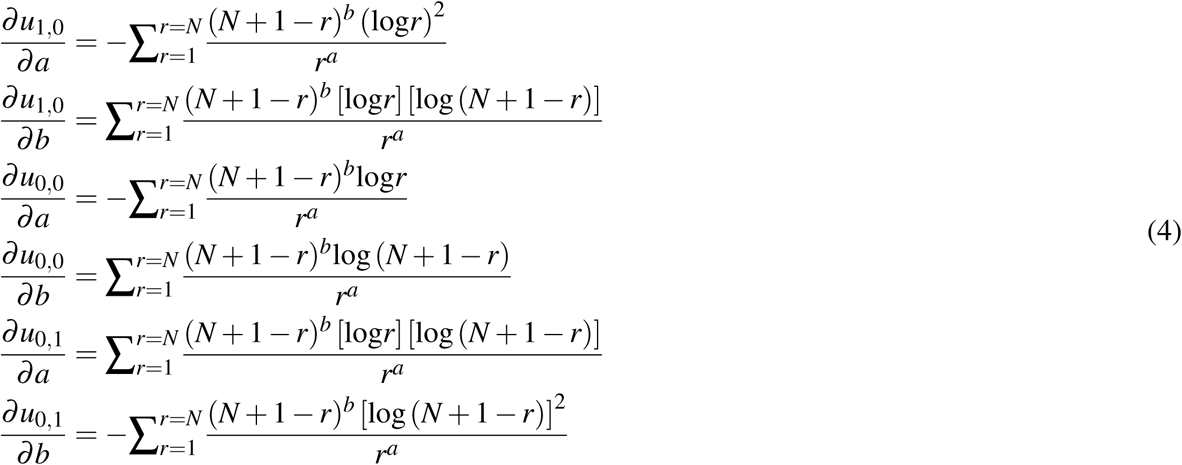

In a compact form, these can be written more generally, for any *i, j* = 0, 1, as

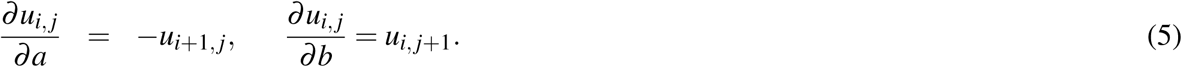

Substituting the above expressions in the formula for Fisher information matrix in Eq (3) of the main paper, we get its simplified form for computation within our ROSeq.

## 2 Extended Result

**Figure 1.**
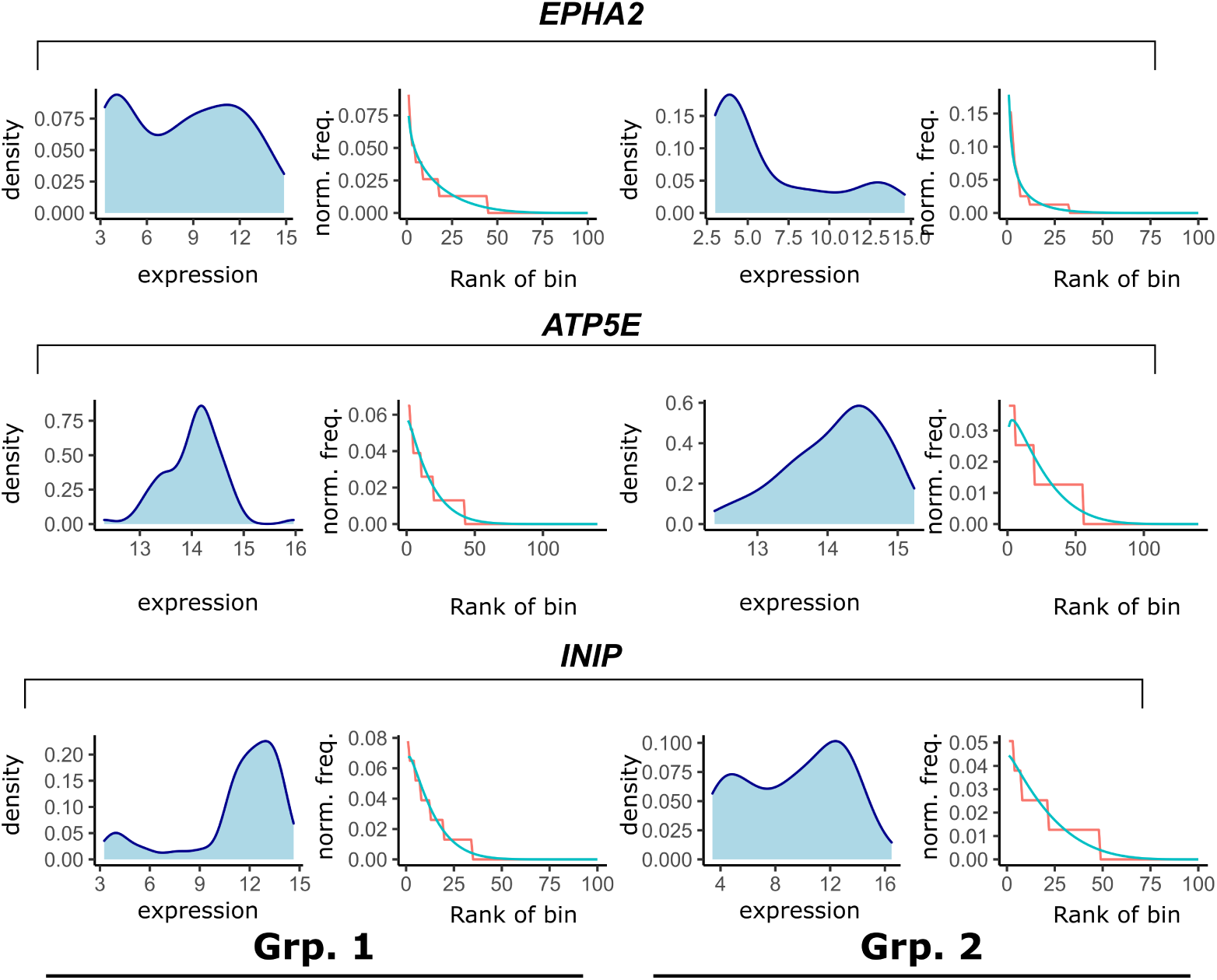
Five randomly selected non-differential genes between myoblasts sampled before/24 hours after differentiation [1]. Differential expression analysis was performed using ROSeq. Each row contains cell-group-wise expression density plot as well as a plot depicting DGBD based modeling of rank-ordered expression bins.

**Figure 2.**
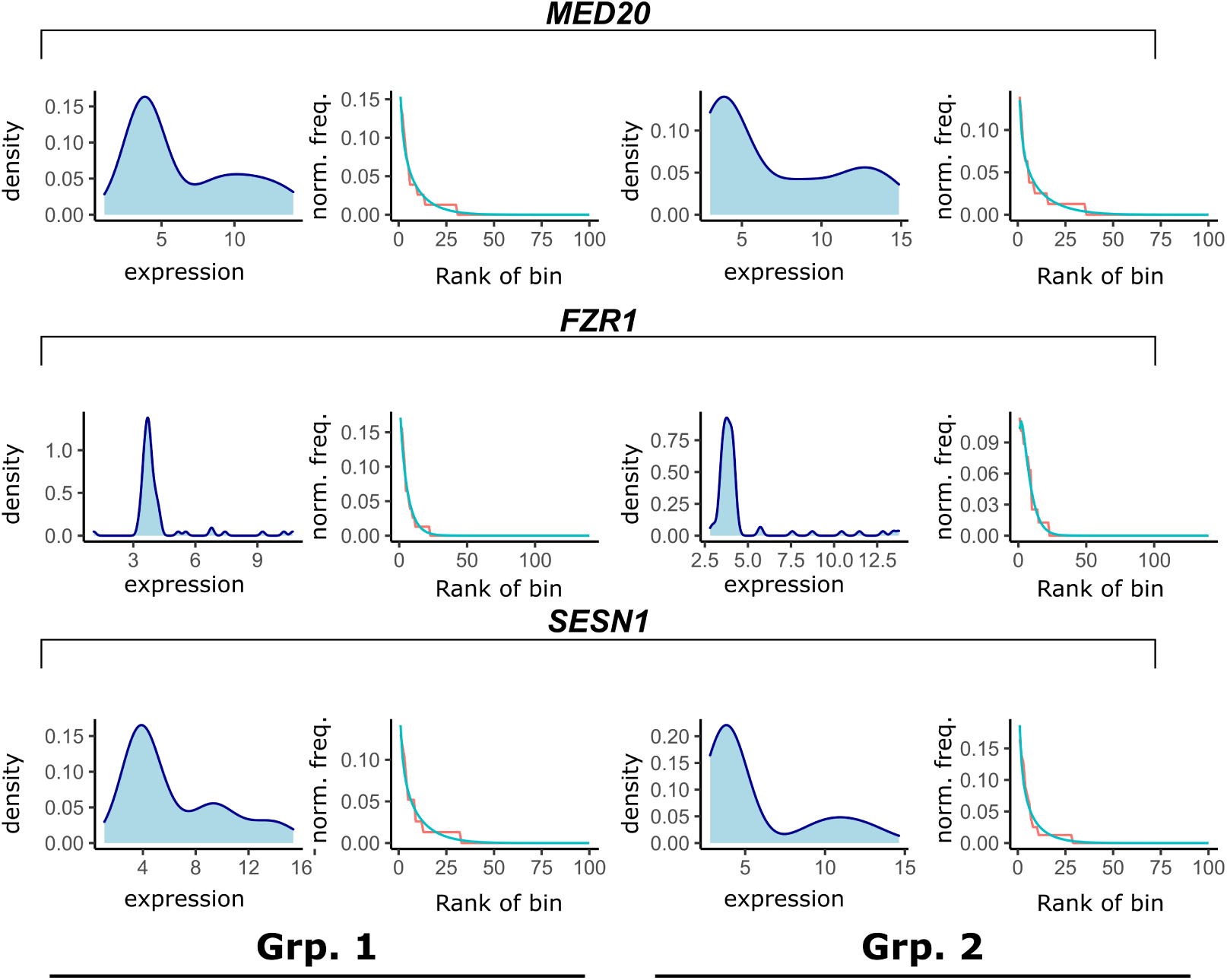
Five randomly selected differential genes between myoblasts sampled before/24 hours after differentiation [1]. Differential expression analysis was performed using ROSeq. Each row contains cell-group-wise expression density plot as well as a plot depicting DGBD based modeling of rank-ordered expression bins.

**Figure 3.**
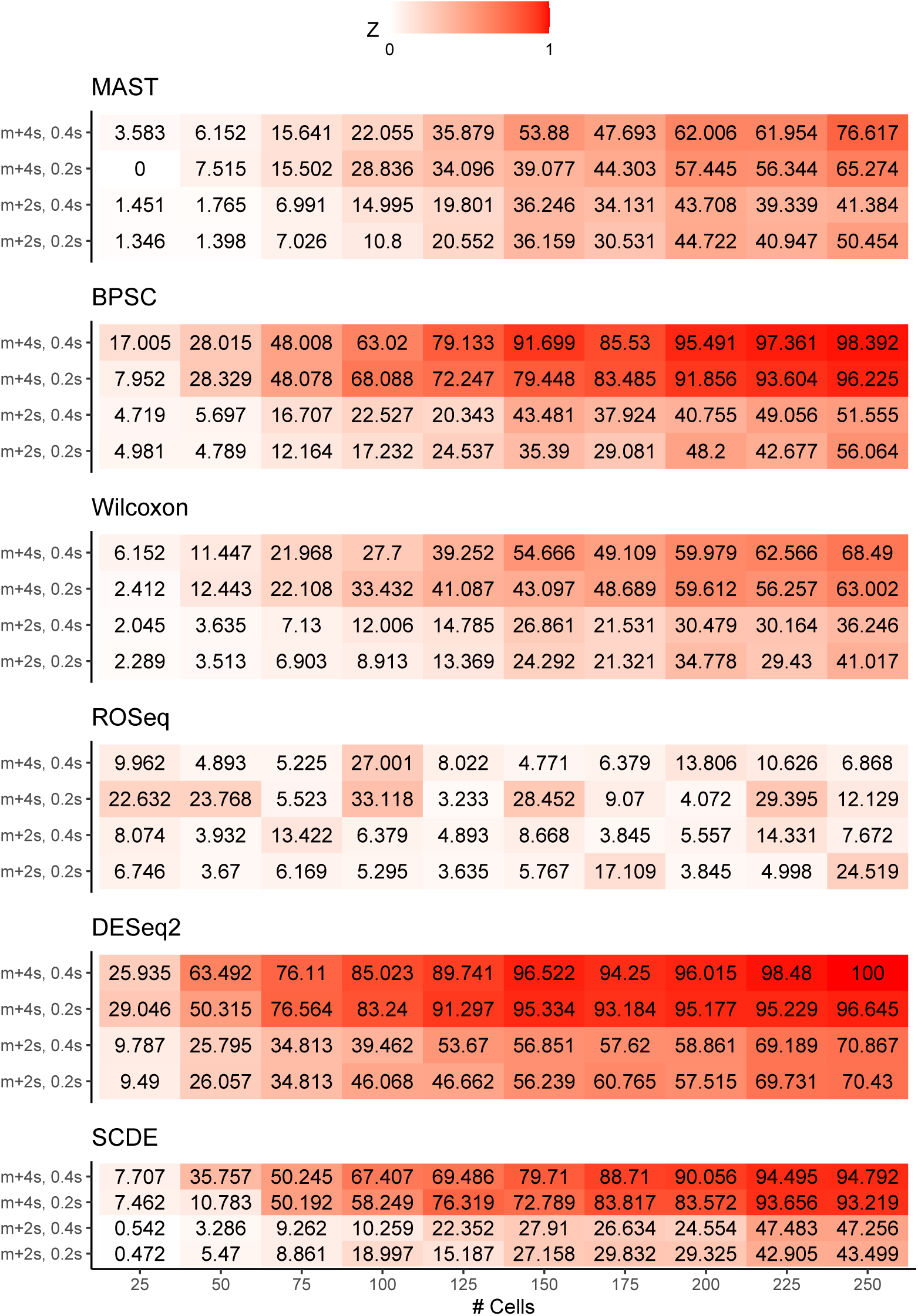
Null datasets were constructed by randomly selecting Jurkat cell transcriptomes into two equal sized groups. The number of cells in the groups varied from 25 to 250, with a step-wise increase of 25. Next, for each gene, one of the cell-groups was chosen, and 20% of the read count values (across cells in the concerned group) were added with simulated Gaussian noise. For a read count value *r* under alteration, the updated read count value *r*′ was set to *r* + *𝒩* (*m* + *c.s, k.s*), where *m* is the mean and *s*, the sample standard deviation of the read count values for the concerned gene across cells of the chosen group. Notably *c* and *k* are constants. We experimented with *c* = 2, 4, and *k* = 0.2, 0.4.

**Figure 4.**
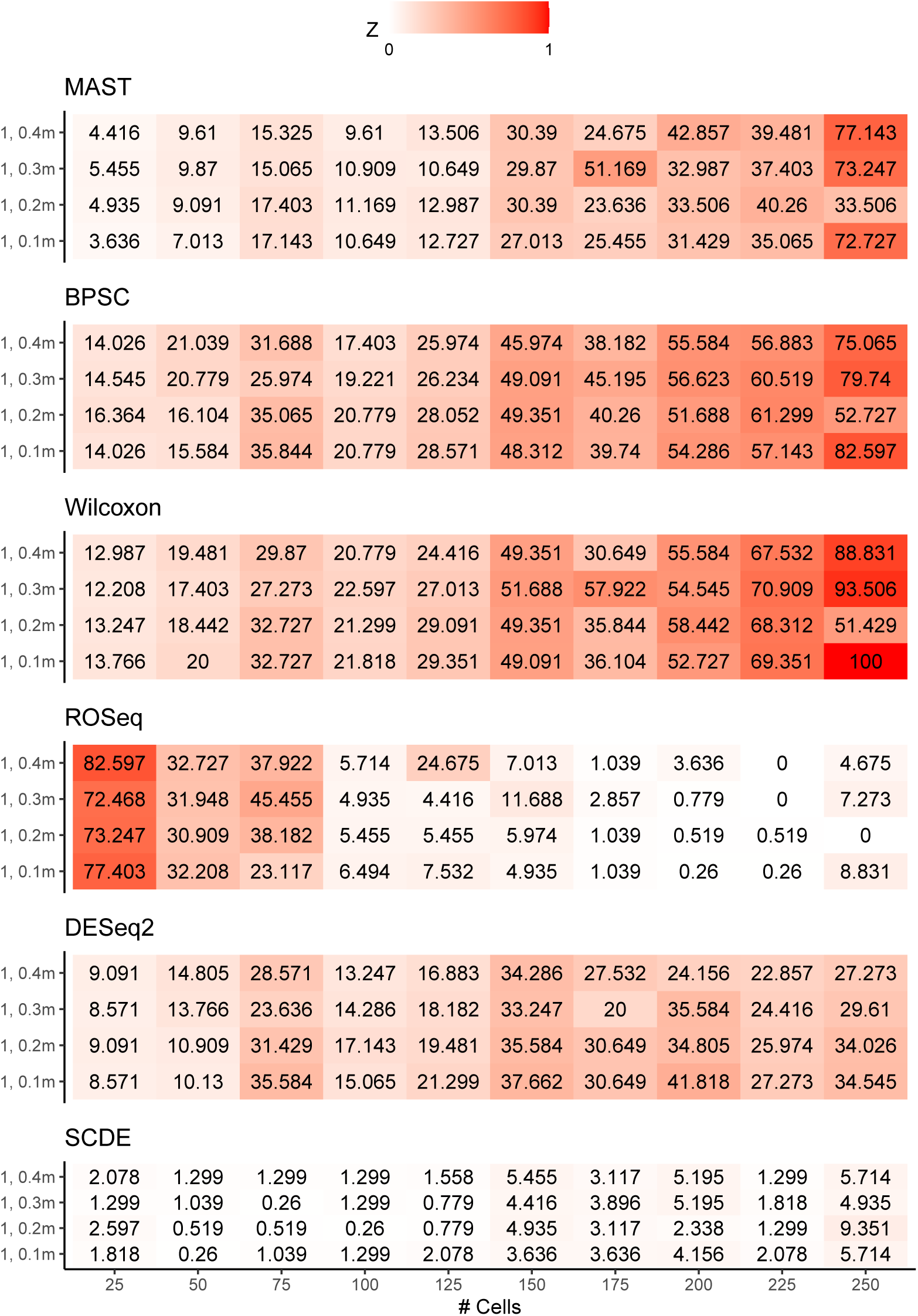
Null datasets were constructed by randomly selecting Jurkat cell transcriptomes into two equal sized groups. The number of cells in the groups varied from 25 to 250, with a step-wise increase of 25. Next, for each gene, one of the cell-groups was chosen, and 10% of the read count values (across cells in the concerned group) were added with simulated uniform noise. For a read count value *r* under alteration, the updated read count value *r*′ was set to *r* + 𝒰 (0, *k.m*), where *m* is the mean of the concerned gene across cells of the chosen group. Notably *k* is a constant. We experimented with *k* = 0.1, 0.2, 0.3, 0.4.

**Figure 5.**
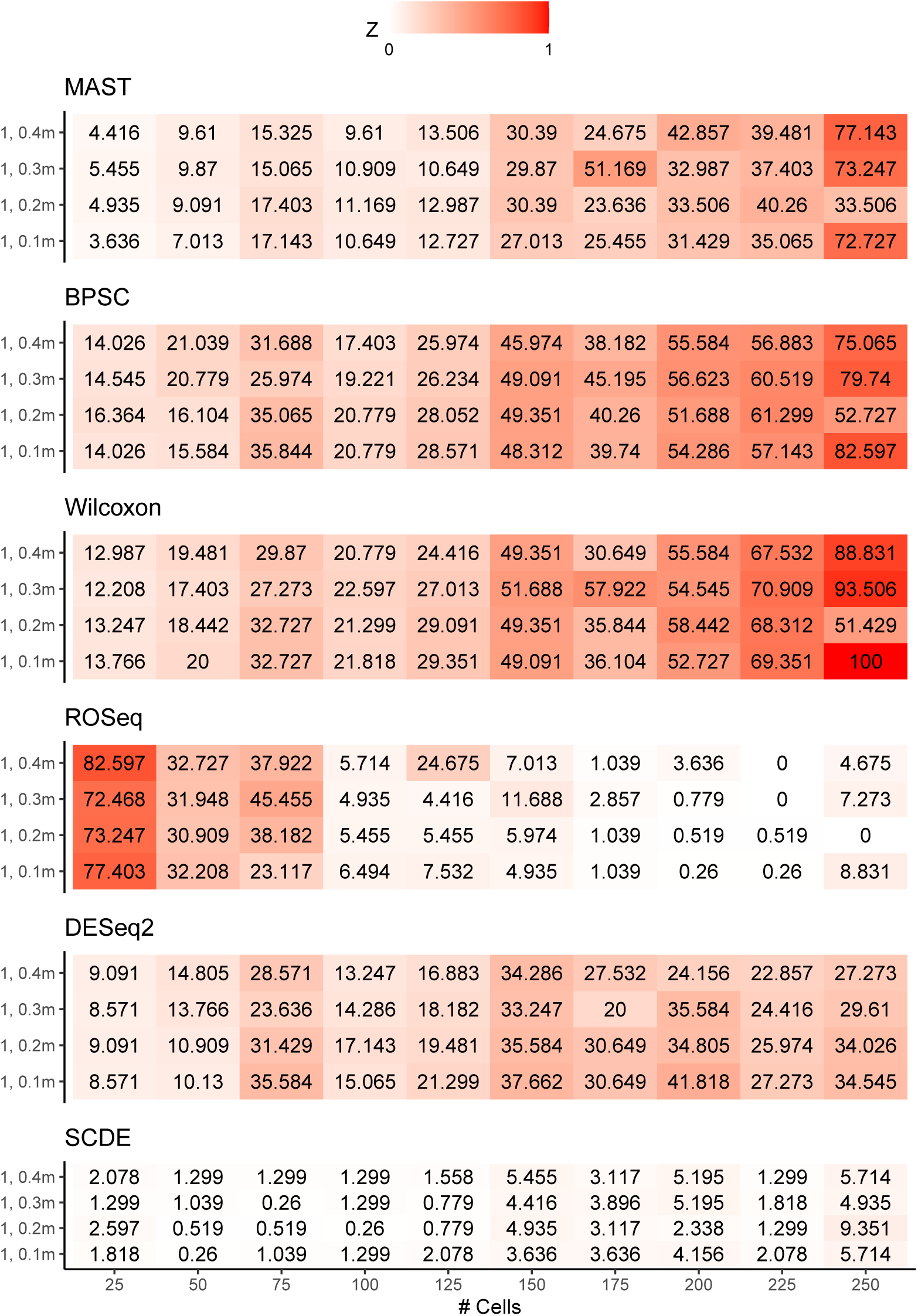
Null datasets were constructed by randomly selecting Jurkat cell transcriptomes into two equal sized groups. The number of cells in the groups varied from 25 to 250, with a step-wise increase of 25. Next, for each gene, one of the cell-groups was chosen, and 10% of the read count values (across cells in the concerned group) were added with simulated uniform noise. For a read count value *r* under alteration, the updated read count value *r*′ was set to *r* + *𝒰* (1, *k.m*), where *m* is the mean, the sample standard deviation of the read count values for the concerned gene across cells of the chosen group. Notably *k* are constants. We experimented with *k* = 0.1,.2,.3,.4.

