## Supplementary Table for "ROSeq: Modeling expression ranks for noise-tolerant differential expression analysis of scRNA-Seq data"

| Data | Method | MCC | F1 | Precision | Recall | Kappa | Accuracy |
|---|---|---|---|---|---|---|---|
| Trapnell data (T0 vs. T24) | MAST | 0.34 | 0.44 | 0.67 | 0.33 | 0.31 | 0.76 |
| | BPSC | 0.46 | 0.54 | 0.61 | 0.49 | 0.46 | 0.85 |
| | Wilcoxon | 0.38 | 0.47 | 0.70 | 0.35 | 0.34 | 0.77 |
| | ROSeq | 0.46 | 0.53 | 0.77 | 0.41 | 0.43 | 0.81 |
| | DESeq2 | 0.46 | 0.54 | 0.57 | 0.52 | 0.46 | 0.86 |
| | SCDE | **0.60** | 0.66 | 0.71 | 0.62 | **0.60** | 0.90 |
| Tung data (NA19098 vs. NA19101) | MAST | 0.04 | 0.01 | 0.98 | 0.01 | 0.00 | 0.29 |
| | BPSC | 0.06 | 0.02 | 0.94 | 0.01 | 0.01 | 0.54 |
| | Wilcoxon | 0.07 | 0.02 | 0.98 | 0.01 | 0.01 | 0.54 |
| | ROSeq | **0.17** | 0.08 | 0.79 | 0.04 | **0.07** | 0.92 |
| | DESeq2 | 0.05 | 0.02 | 0.85 | 0.01 | 0.01 | 0.53 |
| | SCDE | 0.10 | 0.03 | 0.94 | 0.02 | 0.02 | 0.73 |
| Tung data (NA19098 vs. NA19239) | MAST | 0.09 | 0.03 | 0.99 | 0.02 | 0.02 | 0.44 |
| | BPSC | 0.09 | 0.04 | 0.99 | 0.02 | 0.02 | 0.46 |
| | Wilcoxon | 0.09 | 0.04 | 0.99 | 0.02 | 0.02 | 0.47 |
| | ROSeq | **0.32** | 0.21 | 0.94 | 0.12 | **0.19** | 0.93 |
| | DESeq2 | 0.09 | 0.04 | 0.95 | 0.02 | 0.02 | 0.49 |
| | SCDE | 0.14 | 0.06 | 0.98 | 0.03 | 0.04 | 0.67 |
| Tung data (NA19239 vs. NA19101) | MAST | 0.07 | 0.03 | 0.99 | 0.01 | 0.01 | 0.44 |
| | BPSC | 0.08 | 0.03 | 0.99 | 0.01 | 0.01 | 0.48 |
| | Wilcoxon | 0.08 | 0.03 | 0.97 | 0.01 | 0.01 | 0.48 |
| | ROSeq | **0.18** | 0.08 | 0.87 | 0.04 | **0.07** | 0.86 |
| | DESeq2 | 0.06 | 0.03 | 0.87 | 0.01 | 0.01 | 0.50 |
| | SCDE | 0.12 | 0.04 | 0.99 | 0.02 | 0.03 | 0.68 |
